# Rapid histological quantification of muscle fibrosis and lysosomal activity using the HSB colour space

**DOI:** 10.1101/2022.08.02.502489

**Authors:** John C.W. Hildyard, Emma M.A. Foster, Dominic J. Wells, Richard J. Piercy

## Abstract

**Introduction:** Fibrosis is a key feature of many chronic myopathic disorders, such as in the muscle-wasting condition, Duchenne muscular dystrophy. Fibrosis disrupts skeletal muscle architecture, limits muscle function, impairs regeneration and might reduce efficacy of therapeutic interventions: quantifying muscle fibrosis is thus of key value in monitoring disease progression (or response to treatment) in both pre-clinical and clinical settings. Fibrosis can be visualised histologically via staining with picrosirius red, but its quantification can be time consuming and subject to investigator bias: a rapid, reliable and user-friendly means of quantifying muscle fibrosis in histological images is currently lacking.

**Methods:** We investigated whether the Hue/Saturation/Brightness (HSB) colour-space could be used to quantify connective tissue content in picrosirius red (PSR)-stained muscle sections, using multiple healthy and dystrophic muscles, sampled from two animal models of Duchenne muscular dystrophy (the *mdx* mouse and the DE50-MD dog), at different ages.

**Results:** HSB-based analysis allows muscle fibres, connective tissue and slide background to be readily distinguished in PSR images using only a minimal set of parameters, and correctly identifies fibrotic accumulation under conditions where progressive fibrosis is expected. We have developed an imageJ macro that allows semi-automated high-throughput measurement of fibrotic accumulation, and then further extended our method to demonstrate its validity with another histological stain (acid phosphatase) to quantify lysosomal activity.

**Conclusions:** Histological analysis of muscle pathology is challenging and time consuming, especially with large collections of images: our methods permit fibrotic accumulation to be quantified in such collections rapidly and easily in open-source software, with minimal hardware requirements, and the underlying methodology can be readily extended to other colorimetric histopathological stains.

## INTRODUCTION

The architecture of skeletal muscle is organised and hierarchical, and connective tissue plays a key role in this structure, forming distinct extracellular matrix (ECM) environments at each level. Individual myofibres are surrounded by a thin endomysial layer rich in capillaries and nerves; these fibres are arranged into discrete fascicles: bundles of fibres demarked by a thicker and more collagenous perimysial layer; fascicles are themselves bundled within the epimysium, a yet thicker layer of matrix material that bounds individual muscles and extends beyond to form the myotendinous junctions [1]. Muscle is highly regenerative, with a dedicated stem cell population (the satellite cells) able to activate, proliferate and differentiate to repair even severe injuries [2, 3]. This recovery is strongly influenced by the ECM environment: injuries that destroy myofibres while preserving the matrix and its basement membrane surrounding those fibres can be repaired almost entirely, while injuries that disrupt both fibre and ECM networks are resolved less completely [4]. Conditions of chronic damage such as those found in the muscle-wasting disease Duchenne muscular dystrophy (DMD) lead to disruption of ECM architecture and concomitant fibrosis, largely initiated by infiltrating fibroblasts: progressive accumulation of aberrant connective tissue (predominantly collagens I and III) [5-7]. This intramuscular scarring disrupts the stem cell niche (impairing ongoing regeneration) [8], alters the mechanical properties of the muscle (increased stiffness, lower force generation) [9], and can even impede blood supply and slow re-innervation after injury [10]. Muscle fibrosis also represents a potential impediment to drug delivery, either by preferential fibroblast uptake of a given therapeutic, trapping of drug within fibrotic deposits, or by simply acting as a physical barrier surrounding the remaining muscle fibres, preventing drug access [11]. Given the clinical and pre-clinical importance of muscle fibrosis, quantitative measurement is of value both to primary diagnostics and in longer-term monitoring of disease progression (or response to therapeutic intervention).

### Quantifying fibrosis

Fibrotic content can be evaluated in several ways, each addressing different aspects of fibrotic accumulation. Measurement of collagen mRNA (particularly collagens I, III and VI) can detect ongoing fibrotic remodelling, where infiltrating fibroblasts are actively proliferating, but this fares poorly within the more transcriptionally-static environment of established fibrosis: collagen is a phenomenally stable protein [12] (surviving fragments have been detected within fossilized dinosaur bones [13]), and consequently, once produced, turnover is low [14, 15]. Measurement of bulk collagen protein as an alternative, is unfortunately challenging: the macromolecular complexes of collagen fibrils exhibit poor solubility, necessitating specialised extractions (acid, protease digestion [16]) in preparation for western blotting, placing concomitant limits of quantification. The collagen-specific modified amino acid hydroxyproline can be assayed via colorimetric chemistry, though this approach necessitates extensive acid hydrolysis of samples and employs strong oxidising agents [17, 18], rendering it laborious and technically demanding.

Bulk assay methods also offer little information beyond raw collagen content (at mRNA or protein level), while collagen content within a muscle sample can be elevated for entirely physiological reasons (proximity to tendon attachment site, high epimysial content, or even simply a more compact fascicular structure). For these reasons, histological assessment of muscle sections is preferred: allowing the investigator to restrict assessment only to representative regions of tissue.

Immunohistochemical/immunofluorescent (IHC/IF) approaches can be used, labelling one or more collagens (or other extracellular matrix components such as fibronectin or laminin), but availability of appropriate antibodies limits the utility of this approach, especially across multiple species. Histological colorimetric stains are more broadly applicable, and here the preferred stain is picrosirius red (PSR): here, muscle fibres appear a vibrant picric-acid yellow, while the surrounding collagen of connective tissue stains prominently with Sirius red. This stain is more reproducible than trichrome stains [19], and as a permanent colorimetric method it allows slide archiving, making it a favoured choice for histological assessment of fibrosis. Staining is moreover comparatively quick, inexpensive and straightforward to conduct: though traditionally restricted to paraffin-embedded sections, a simple xylene pre-treatment [20] renders fresh-frozen sections (preferred in the muscle field) PSR-compatible.

### Quantifying PSR-stained skeletal muscle

Quantification of muscle fibrosis via PSR staining is not straightforward, partly due to the lack of a clearly defined threshold above which connective tissue should be considered pathological. This stain is more commonly used to evaluate fibrosis in non-muscle tissues (liver, kidney, lung, skin [21]), each of which differ in tissue architecture, and each of which consequently exhibit distinct distributions of pathological fibrosis, none of which readily translate to muscle. PSR staining can be visualised via polarized light, fluorescence, or conventional brightfield microscopy, but at present there is no standard method of quantifying skeletal muscle fibrosis: most muscle studies concern cardiac tissue (and even here debate remains over the most effective approach [22]). Stereology using polarizing microscopy is considered the ‘gold standard’ for measuring cardiac fibrosis [20, 23]: PSR staining enhances the birefringence of collagen fibres, and when viewed through a polarizing filter only collagen content is resolved, offering a rapid method for quantifying total fibrosis. This translates well to skeletal muscle (Fig 1A), but as birefringence exploits *en bloc* properties of collagen fibres, smaller deposits of connective tissue (such as endomysium) are resolved less readily, if at all, and distinction between non-collagen tissue and slide background is impossible. Extent of polarization also correlates with the precise alignment (and type) of collagen fibres [21], and signal intensity can consequently alter with sample orientation [24], necessitating averaging of images taken at multiple orientations (or use of circularly-polarized, rather than plane-polarized light). Access to a polarizing microscope is essential, and stereological reconstruction is also time-consuming and comparatively specialized, limiting widespread utility.

**Figure 1:**
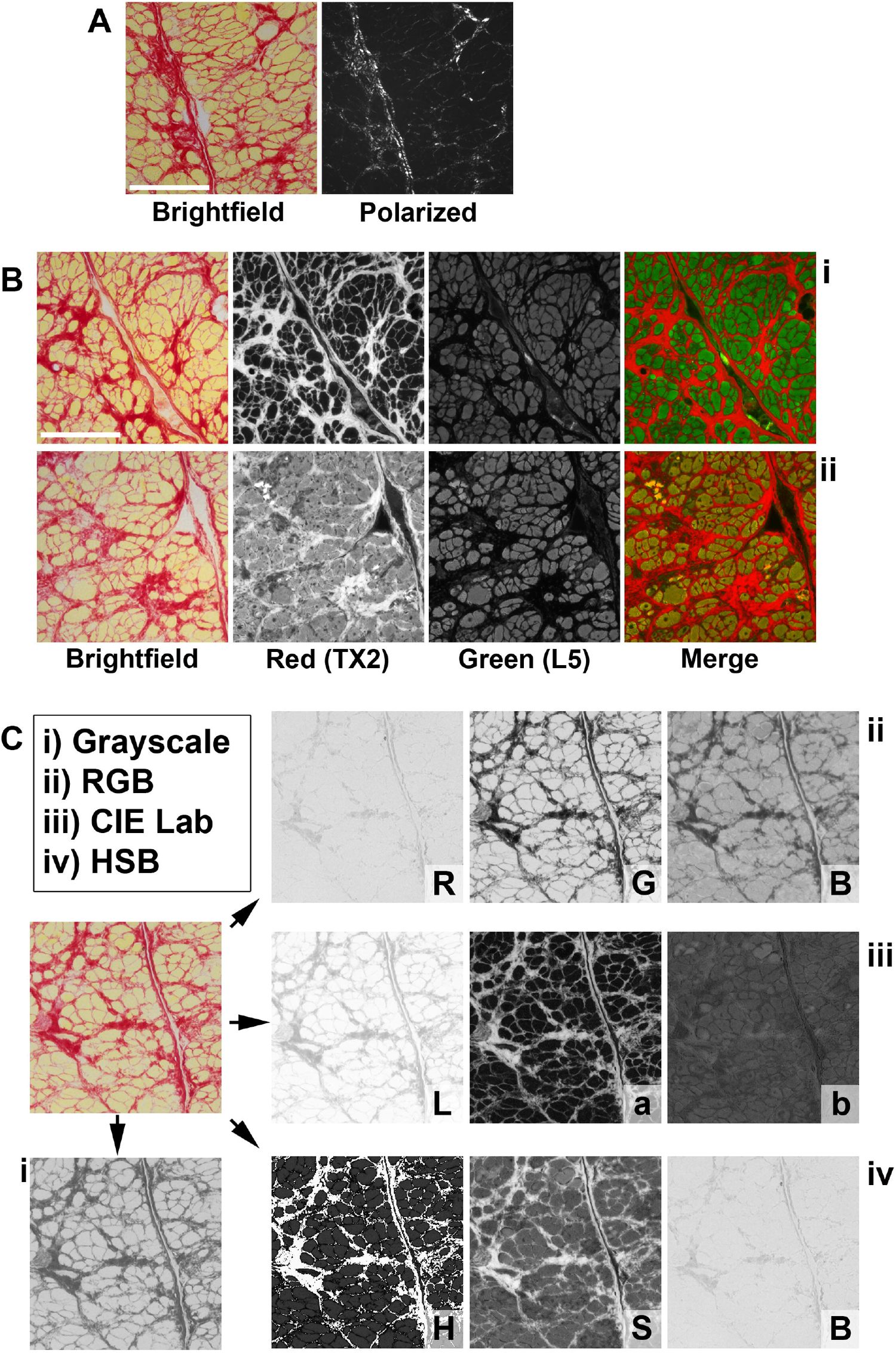
Histological analysis of PSR-stained muscle sections Polarized light microscopy (A) shows birefringence of collagen fibrils in PSR-stained muscle sections, though thinner connective tissue regions (such as endomysium) are poorly resolved. Fluorescence microscopy (B) exploits the red fluorescence of Sirius red-stained connective tissue (TX2 filter -texas red/alexafluor 594) and the green fluorescence of picric acid-stained muscle fibres (L5 filter - FITC/GFP/alexfluor 488), though samples (comparable under brightfield) can exhibit substantial red-fluorescence variability under (B i vs B ii). Brightfield microscopy images (C) can be segregated simply via conversion to grayscale, with loss of all colour information (i), or split into red/green/blue (RGB) channels (ii), Lightness/a/b (Lab) channels (iii) or hue/saturation/brightness (HSB) channels, with variable utility offered by each. Scalebars: 200µm.

An alternative approach is fluorescence microscopy: both picric acid and Sirius red exhibit fluorescence spectra with separable characteristics [24, 25]: yellow-stained muscle fibres fluoresce at ‘green’ excitations (Alexa fluor 488, FITC, GFP), red connective tissue at ‘red’ (Alexa fluor 594, Texas red) (Fig 1B). Separation of muscle and connective tissue into distinct channels allows these elements to be evaluated in isolation. This method naturally requires a fluorescent microscope, and imaging in multiple channels increases user-time needed for data collection considerably. In our experience this method is also particularly subject to substantial batch-to-batch variation: stains that appear wholly comparable under light microscopy can show significant intrafibrillar Sirius red signal via fluorescent methods, rendering them challenging or even impossible to quantitatively compare (Figure 1B, i vs ii).

Brightfield microscopy offers most utility and accessibility, and there is consequently considerable interest in quantifying images obtained via this approach (though at present, no clear consensus on a suitable method). Conversion of a colour image to greyscale (or monochrome white/black) permits relatively straightforward densitometry, as the Sirius red component naturally maps to darker shades (Figure 1C, i). The advantages offered by the dual-colour nature of the stain are however lost, precluding identification of slide background. RGB-based analyses (Red/Green/Blue) perform poorly, as red and yellow components share considerable overlap when expressed as RGB values (Figure 1C, ii). Differences in stain intensity and uneven illumination can further confound efforts to use this method quantitatively. Use of the CIE ‘Lab’ colour-space [26] was proposed by Hadi *et al* [20]: this defines pixels by 8-bit luminance (L), along with ‘a’ and ‘b’ channels (representing green/red balance and blue/yellow balance respectively). While not immediately intuitive, yellow and red components are segregated effectively via this approach (Figure 1C, iii), and accordingly, Hadi *et al* developed a semi-automated method for image quantification, using a nested series of empirically-determined thresholding filters to isolate regions denoted as ‘lumen’, ‘fibrosis’ or ‘cardiomyocytes’, a method shown to be comparable to stereology while accelerating analysis time and accessibility [20]. As implied by the category names, however, this method was developed for cardiac tissue, using fixed thresholds appropriate for rat heart (the authors subsequently assessed liver, kidney and lung, but skeletal muscle was not evaluated).

To develop a more intuitive method (and one specifically optimised for skeletal muscle), we have investigated use of the hue/saturation/brightness (HSB) colour-space. Like RGB and Lab colour-spaces, HSB describes each pixel of an image using three 8-bit (0-255) values, but here these three values denote hue, saturation, and brightness (Figure 1C, iv). A key advantage of this colour-space is that the entire colour spectrum is mapped to a single channel (hue): saturation and brightness channels indicate intensity and illumination, but a given hue value remains constant regardless of these factors. As shown in Figure 2, changes in saturation or brightness (reflecting stain intensity/section thickness and microscope illumination, respectively) can significantly alter RGB or Lab values, and do so in a counter-intuitive fashion that is challenging to address quantitatively. Conversely, under HSB assignment, red hues retain their specific values (and remain distinct from orange or yellow values) even if heavily desaturated or darkened.

**Figure 2:**
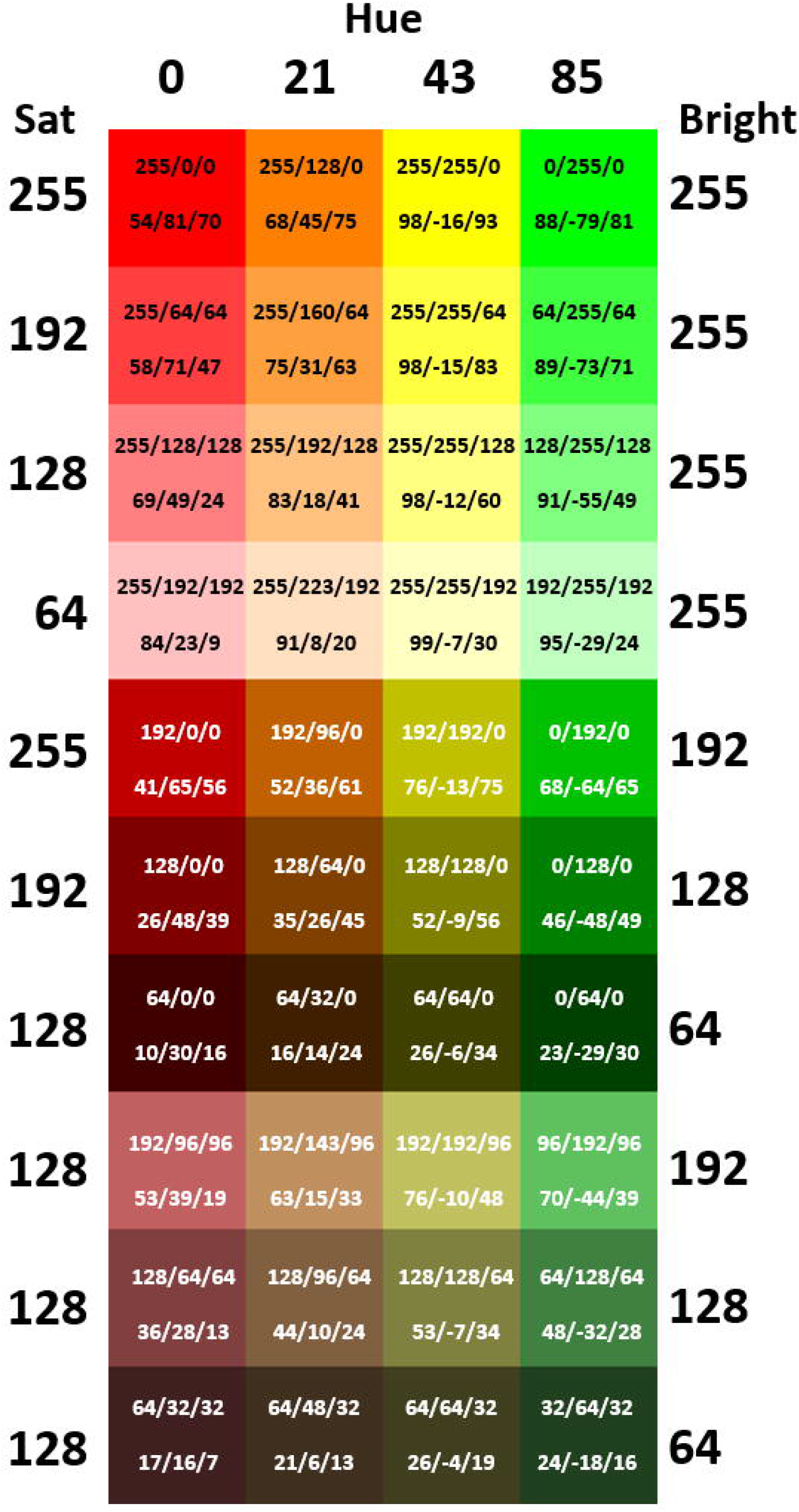
The HSB colour-space renders colour independent of brightness or saturation The top row depicts shades of red, orange, yellow and green. The same colours are then shown with decreasing saturation (rows 2-4), decreasing brightness (rows 5-7), or both (rows 8-10). Hue, saturation and brightness values are displayed around the periphery, with corresponding RGB (upper) and Lab (lower) assignments shown within each square. Hue values remain unchanged regardless of saturation (corresponding to stain intensity/section thickness) or brightness (slide illumination), while both RGB and Lab assignments vary substantially with such changes.

Our data shows that this space is well suited to analysis of PSR-stained histological images, and moreover this approach can be adapted to a high-throughput semi-automated macro format while retaining investigator flexibility. We have validated our method using muscle sections taken from healthy and dystrophic (*mdx*) mice, and healthy and dystrophic (DE50-MD) dogs, showing that this method is sensitive, reproducible, highly robust, and of broad utility. We further show that the HSB colour space is sufficiently versatile to allow the same principal method to be adapted to other stains of pathological interest, such as acid phosphatase, a commonly used stain that identifies tissue lysosomal activity encountered in (for example) inflammatory conditions.

## METHODS

### Sample collection

All samples used in this study were taken from archived collections: no animals were sacrificed specifically for this work. As described previously [27-29], samples were collected as follows:

#### Mouse muscle

mice (*mdx* or healthy strain-matched C57Bl/10) of the indicated ages were killed by cervical dislocation and muscle tissues were harvested immediately. Muscles were mounted in cryoMbed (Bright) on cork blocks and snap-frozen under liquid nitrogen-cooled isopentane before storage at -80°C.

#### Canine muscle

Skeletal muscle samples (∼0.5 cm^3^) were biopsied at 3, 6 and 9 months of age from WT and deltaE50-MD (DE50-MD) dogs by open approach, from the left *vastus lateralis* muscle, with dogs under general anaesthesia as described in [27] as a component of an ongoing natural history trial (Wellcome Trust grant 101550). Samples from a total of six animals (3 WT, 3 DE50-MD) were used for this work. Biopsy samples were frozen and sectioned as for mouse tissue (above).

### Sectioning

All muscles were cryosectioned using an OTF5000 (Bright) at 8µm (dog) or 12µm (mouse) thickness and mounted onto glass slides (Superfrost Plus). Slides were air dried for 30 mins at room temperature and stored in sealed containers at -80°C until use.

### PSR staining

Picrosirius red staining is typically used with paraffin-embedded sections rather than with the fresh frozen sections more commonly used in muscle histology. Post-hoc fixation and dehydration can render frozen tissues compatible with this stain [17], however a simpler method was identified by Hadi *et al* [20], here replicated in full for reference (see also protocols.io version [30]). Steps we have found to be critical are indicated with an asterisk*.

Slides were removed from -80°C storage and allowed to dry and equilibrate to room temperature (30 mins) before being immersed completely in xylene for > 20 mins* (no negative effects were observed with incubations of up to an hour, but 20 min is the minimum). Slides were allowed to dry completely before rehydration through graded alcohols (100%, 90%, 80%, 70%), allowing 3-5 mins in each solution. After a final incubation in water (∼1 min), slides were immersed in picrosirius red solution (0.1% Sirius red F3B in a saturated ∼1.5% solution of picric acid –this stain can be reused multiple times and remains stable for at least 1 year) for 45-50 minutes*. Incubations shorter than 45 minutes result in weaker, washed-out staining, while incubations longer than 50 minutes result in progressive Sirius red staining of muscle fibres. Slides were then washed briefly (<5 sec) in water and incubated in a 0.5% solution of acetic acid (in distilled water) for 2 minutes to fix the stain. Finally, slides were dehydrated through graded alcohols (∼1 minute in 70%, 80%, 2-5 minutes in 90% and 100%), allowed to dry, equilibrated with xylene (1 hour to overnight) and mounted in DPX mounting medium.

### Acid phosphatase staining

Acid phosphatase staining historically involves barbital acetate-buffered solutions: as a (legislature) controlled substance, this is sub-optimal for routine use. We have instead used a sodium acetate-based protocol [31] that gives comparable results, reproduced here in full.

Slides were removed from storage and allowed to equilibrate to room temperature as above, while staining solutions were prepared. For staining solution 1, 200 ml sodium acetate buffer (320mM, pH 4.8) was combined with 4 mg napthol AS-B1 phosphate (1% stock in dimethylformamide-can be stored at -80°C) and mixed well. Separately, staining solution 2 was prepared by mixing 3.2 ml pararosaniline-HCl solution (4% stock in 20% conc. HCl -can be stored in aliquots at -80°C) in dropwise fashion with 3.2 ml sodium nitrite (4% stock in distilled water, prepared fresh) until a straw-coloured solution was produced. This was incubated at room temperature for 2 mins, then added to staining solution 1 and mixed well. pH was adjusted back to 4.8 by careful (dropwise) addition of 1M HCl. Slides were then immersed in staining solution and incubated at 37°C for 2 hours. To terminate the reaction, slides were washed well in distilled water before counterstaining for 1 min in Mayer’s haemalum (Sigma) followed by bluing under running tap water for 3 min. Slides were dehydrated rapidly through graded alcohols (∼30 sec per stage) then allowed to dry completely before equilibration to xylene and mounting in DPX as above.

### Imaging

Images were collected using a DM4000 upright brightfield microscope using a DC500 colour camera (Leica Microsystems), with multiple representative .tif format images at 10x objective (10x HC PL FLUOTAR PH1, NA=0.3) collected per sample. For mouse samples, 5-6 images were collected in most cases, while for the smaller canine muscle biopsy samples, typically 3-5 images were collected (our data suggests that 3-5 images allows representative quantification, but given the ease of image collection, and the speed of our analysis method, we see little disadvantage in collecting additional fields if tissue samples permit it).

### Analysis

All analysis used the Fiji distribution of ImageJ (https://fiji.sc/). Initial investigations employed the ‘colour threshold’ tool, which works within the HSB colour space.

For PSR quantification, the saturation channel was used to exclude background, with remaining pixels then segregated by hue into yellow, red and ‘other’. Pixel counts corresponding to each category were exported. Analysis methods were subsequently incorporated into a batch-compatible imageJ macro (PSR_quantify.ijm [32]) file, which can be executed in Fiji via the plugins>>macro>>run command (see figure 7 for a schematic of the process, and protocols.io [30] for a step-by-step guide). All processing uses image duplicates, leaving source images unchanged (analysis can be repeated with different parameters on the same image repositories without risk). The user provides the macro with a source folder (containing images) and a destination folder (for analysis): the macro then iterates through all compatible images. Analysis occurs in two stages. First, images are converted to HSB stacks, then split into the three corresponding 8-bit greyscale images for hue, saturation and brightness. Histogram data for saturation and hue are extracted, and mean values (+/-standard deviation) of each, for the entire image set, are plotted. Default values for background subtraction (saturation) and red/yellow boundaries (hue) are overlaid. The investigator can accept these default values, or adjust accordingly. Mean/SD and raw histogram values per image are automatically saved as comma separated value files (.csv) for archiving/QC purposes. Analysis occurs in the second stage: for each image, saturation values are used to convert the saturation channel to a background subtraction mask, which is then applied to the hue channel. Using the red/yellow boundary values, pixels in this hue channel are assigned to ‘yellow’, ‘red’, ‘background’ or ‘other’. The ‘fibrosis fraction’ (red pixels as a fraction of total tissue pixels) is then derived (red/red+yellow), effectively representing each image as a value from 0-1. Values are exported and saved automatically (again as .csv files), and the threshold values used are appended to the exported data for future reference.

For acid phosphatase (AP_quantify.ijm [33]), a similar approach is taken: brightness is used to establish background subtraction, with hue used to delineate the ‘red’ boundary, which is further gated based on saturation to restrict quantitation only to stain above a specified intensity (as this stain exhibits modest labelling of all tissue). Following selection of thresholds, images are analysed: the brightness channel is used to generate a mask which is then applied to the hue channel. All non-background pixels are counted, and then a further mask (generated from the saturation channel) is applied to exclude all low saturation pixels. All remaining pixels falling within the selected ‘red’ hue range are then counted and expressed as a fraction of total (the AP fraction). All AP fraction values are then log transformed for analysis.

### Statistics

Per-image fibrosis fraction values were used to general per-sample means: all statistical analysis used these mean per-sample values. Per-image AP fraction values were log transformed, and statistical analysis used the mean of the per-sample log values.

For canine muscles, the repeat biopsy sampling allowed repeated measures 2-way ANOVA (with individual animals as subject, and genotype, age and genotype/age interaction as factors). Post-hoc Holm-Sidak multiple comparisons were used to compare healthy and dystrophic cohorts within each age category. All statistical analysis was conducted using GraphPad Prism 9.4.0 and differences were considered statistically significant when p<0.05.

## RESULTS

### Saturation and hue allow background subtraction and image segmentation

Using muscle sections from the diaphragms of 24-week-old healthy (WT) and dystrophic (*mdx*) mice, we investigated HSB-based image segmentation. Diaphragms were selected for two principal reasons: firstly, fibrosis in the *mdx* diaphragm is profound (other muscles in this animal model exhibit only mild fibrotic changes [34-36]) and secondly, images of diaphragms also typically feature significant areas of slide background, making development of a background subtraction process essential. All images used for analysis were collected using a 10x microscope objective, which we found to be most representative. At lower magnification (5x), endomysial layers surrounding individual fibres are of sub-pixel thickness (and hence poorly resolved), while at higher magnification (20x or greater), collecting representative images is challenging: perimysial or epimysial layers cannot be captured without these regions dominating the imaging field (resulting in high per-image variation). As shown in figure 3, the ‘colour threshold’ tool (which uses HSB by default) readily facilitated both exclusion of unstained slide background and segmentation of stained tissue. Saturation histograms revealed two clear peaks, one corresponding to unstained (low saturation) slide background, the other to stained tissue area. Thresholding via saturation thus restricted analysis only to stained tissue area (figure 3A), and further thresholding based on hue allowed delineation of yellow and red fractions (and extraction of pixel areas corresponding to both: figure 3B). Note that the hue spectrum is technically continuous (i.e. circular) but under 8-bit representation this spectrum is necessarily split for mapping to a 0-255 range. Red hues span the split, and thus map both to the upper (>230) and lower ranges (<20). No thresholding via brightness was required (i.e. segmentation was robust to differences in slide illumination).

**Figure 3:**
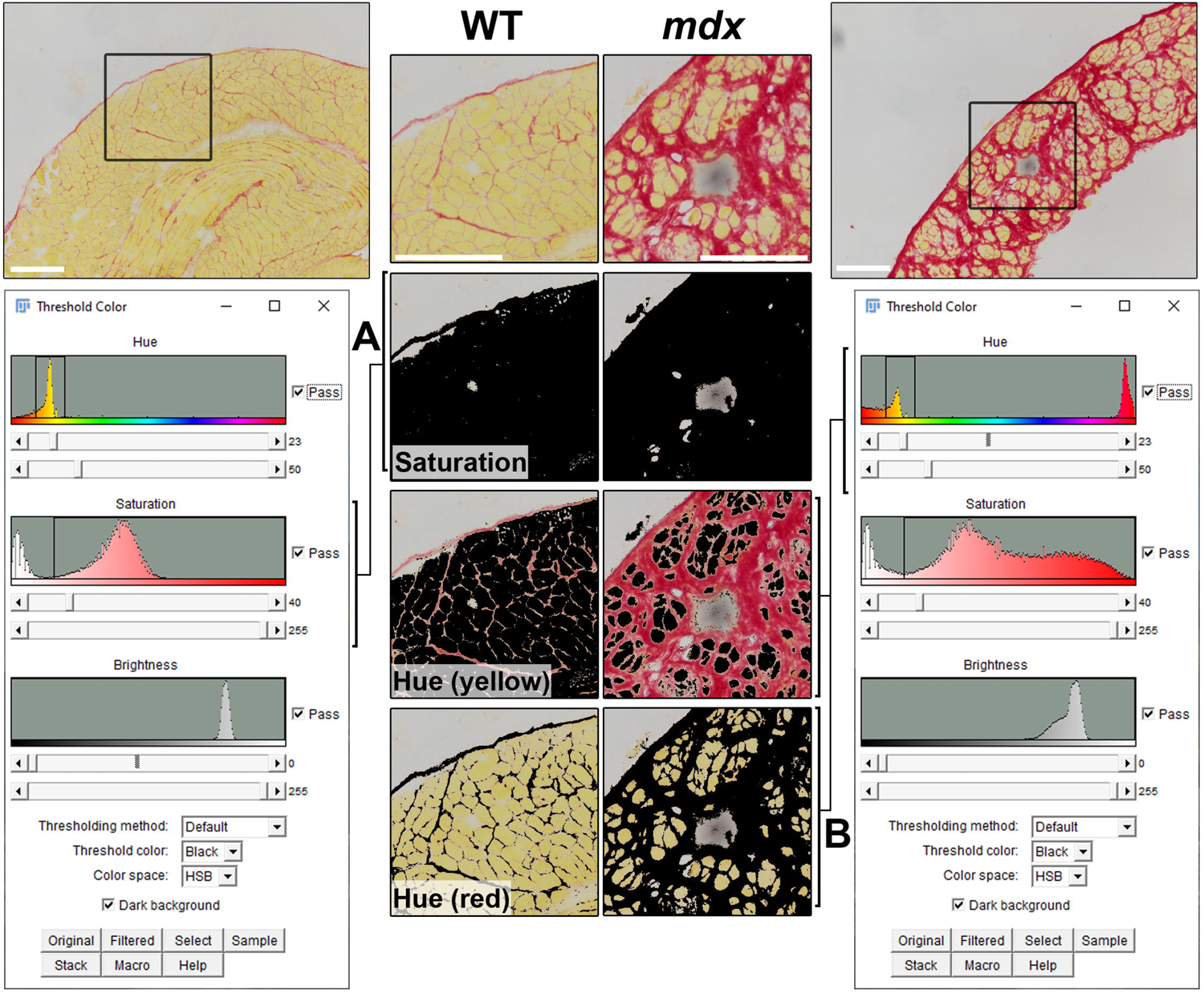
The HSB colour-space permits background elimination and segregation of Sirius red- and picric acid (yellow)-stained tissue regions The HSB-based colour threshold tool in ImageJ permits background pixels to be eliminated via saturation-gating (A), while Hue can be used to segregate remaining pixels into ‘yellow’ and ‘red’ fractions (B). The brightness channel is not required. Both healthy (WT) and dystrophic (*mdx*) samples can use identical settings, avoiding investigator bias. Scalebars: 200µm.

Comparison of multiple images revealed highly consistent hue spectra (figure 4A, B), suggesting identical thresholding values could be applied to entire image collections (whether healthy or dystrophic), eliminating any requirement to manually assign values on a per image basis (both time consuming and crucially, potentially subject to investigator bias). As shown (figure 4B), hue values of low-saturation background pixels generated periodic ‘spikes’ of noise in the histogram, but these were eliminated following subtraction of slide background (figure 4C). Hue values between 23 and 50 were designated ‘yellow’ (picric acid) and hue values of 0-23 and 230-255 were designated ‘red’ (Sirius red): total pixel counts within these brackets (effectively, area under the curve: representing a fraction of total imaging field pixels) were extracted for each image and converted to a single ‘fibrosis fraction’ value by expressing ‘red’ as a fraction of ‘total’ (red + yellow). Per-image values were in good agreement, particularly for dystrophic muscle, suggesting that this metric is representative of a given sample, and 3-5 images per section is sufficient for quantification (figure 4D).

**Figure 4:**
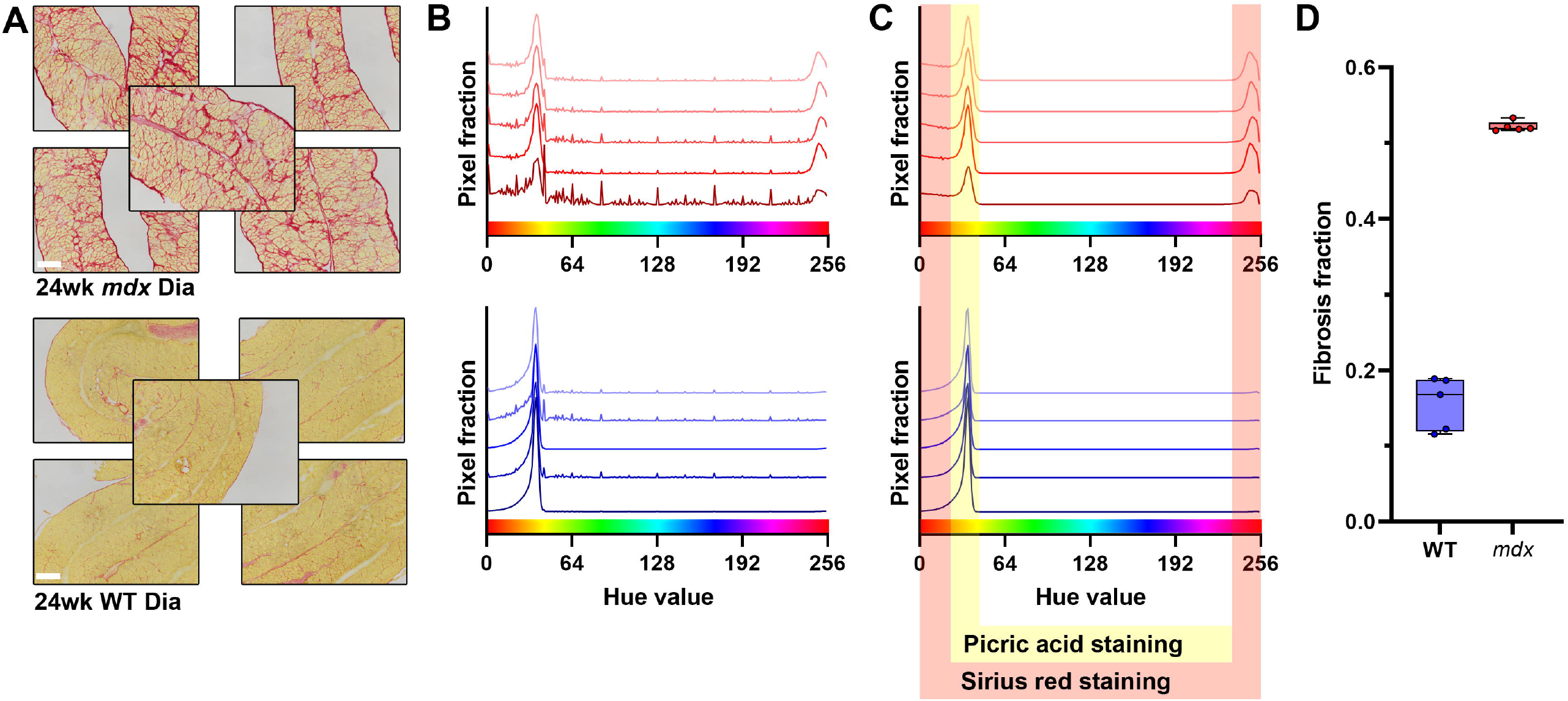
Hue spectra are highly consistent between images, and allow image quantification (A) 5 representative images collected from 24-week-old *mdx* diaphragm (top) and 24-week-old WT diaphragm (bottom). Raw hue spectra are consistent between images and between genotypes (B), with periodic spikes corresponding to stochastic hue assignment of low-saturation background pixels. Hue spectra after background subtraction (C) remain consistent and allow area (pixel counts) of Sirius red- and picric acid-stained tissue to be extracted and used to generate a quantitative ‘fibrosis fraction’ value (red pixels/(red+yellow pixels)). Per image fibrosis fraction values are highly consistent within a given sample (D) and readily permit quantification of fibrosis in healthy and dystrophic muscle. Scalebars: 200µm.

### Fibrosis fraction is a robust quantitative parameter

To assess the quantitative utility of this analysis approach, we investigated a larger cohort of *mdx* and WT mouse samples of different ages (from our archive collection, stained on separate occasions). Two different muscles were measured: the diaphragm and the tibialis anterior (TA) muscle. The TA is one of the more widely studied muscles in the *mdx* mouse, but does not exhibit marked fibrotic accumulation (as noted above [34-36], fibrosis in the *mdx* mouse is essentially a diaphragm-specific phenomenon). Using diaphragm and TA samples thus tests the upper and lower bounds of a quantitative approach. We further ensured that multiple staining batches were included, as effective image quantification should be robust to batch-to-batch variation. As with the traces shown in figure 4B, hue spectra suggested high consistency both between muscle type and staining batch, thus identical thresholding parameters were applied to all samples. Derived per-image fibrosis fractions showed good agreement within each sample and mean per-sample values were also highly consistent by age and genotype (figure 5A), revealing progressive fibrotic accumulation within dystrophic, but not healthy diaphragms. Conversely, both healthy and dystrophic TA muscles (figure 5B) showed no cumulative fibrosis over the ages measured (though note higher values in 6-week-old *mdx* samples, reflecting Sirius red staining of calcified fibres that accompany the widespread necrosis at this age). The strict red/yellow thresholding of this approach is not without potential caveats: while the hue value at which the ‘yellow’ range ends (∼50), and the point at which the upper portion of the ‘red’ range begins (∼230), are both trivial to determine (see figure 4C), this strict red/yellow thresholding also necessitates a defined boundary between these two hues (no pixels are assigned as ‘orange’). High boundary values will necessarily increase ‘red’ fraction while decreasing ‘yellow’ (and vice versa), potentially placing significant weight on this value. A boundary of 23 was used for the analysis shown in figure 5A and B: accordingly, to establish the robustness of our quantification, we reanalysed diaphragm samples using higher (25) or lower (20) threshold values for the red/yellow boundary, essentially measuring the sensitivity of this parameter. As shown (figure 5C i-iii), marginally higher/lower derived fibrosis fractions were obtained as expected, but this occurred in an *en bloc* manner, leaving the magnitude of the difference between ages/tissues essentially unchanged. Precise selection of boundary value thus does not appear to be critical, provided the same value is applied consistently.

**Figure 5:**
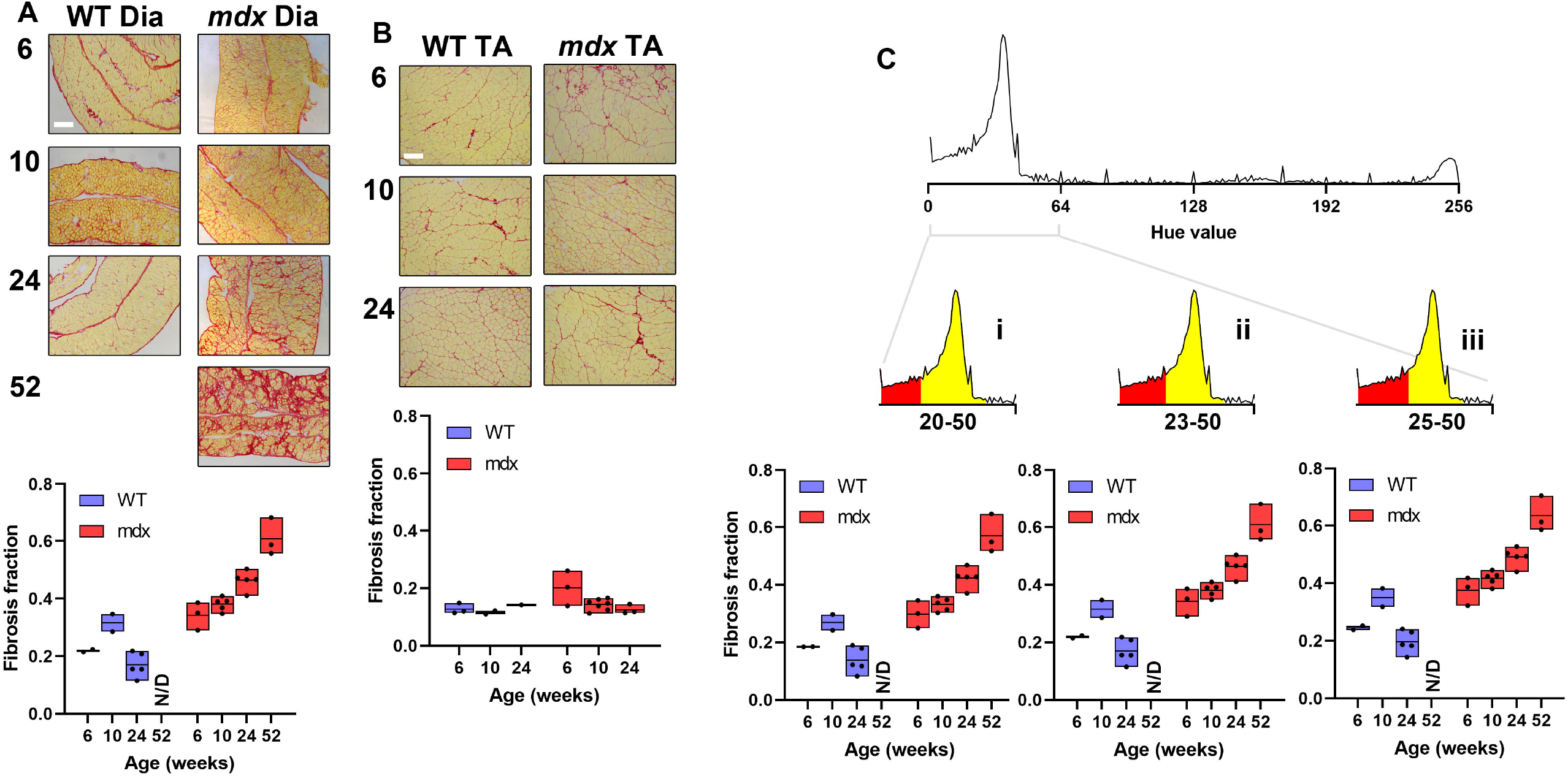
HSB-based fibrosis fraction is a robust quantitative metric Quantitative image analysis of mouse diaphragm muscle (A) at 6, 10, 24 and 52 weeks of age (as indicated) reveals clear age-associated increases in fibrosis fraction within dystrophic (mdx), but not healthy (WT) animals. In tibialis anterior muscle (B) at 6, 10 and 25 weeks of age, fibrosis factions show no age-associated changes whether healthy (WT) or dystrophic (mdx). Fibrosis fraction values are largely robust to selection of red/yellow boundary (C): analysis of diaphragm data using a lower (i), intermediate (ii) or higher (iii) hue boundary results in downward/upward shifts in measured fibrosis fraction values but retains differences between groups. Scalebars: 200µm.

As noted above, dystrophic mouse muscle represents a test of extremes: fibrosis in the diaphragm is profound while fibrosis in the TA is essentially absent. To test the utility of this approach over more intermediate states of fibrotic pathology, we assessed muscle samples taken from the DE50-MD canine model of DMD. Dystrophic dogs exhibit systemic, progressive skeletal muscle fibrosis similar to human patients [37-41], and the greater muscle bulk of dogs (compared to mice) allows repeat biopsy, permitting fibrosis to be monitored within the same muscle over time. Image analysis of *vastus lateralis* muscle samples biopsied from 3 DE50-MD dogs and 3 WT littermates at 3, 6 and 9 months (figure 6) detected elevated levels of connective tissue in dystrophic muscle overall (P<0.01, DE50 vs WT), and between matched ages following multiple comparisons (P<0.05 in all cases). Sensitivity was moreover sufficient to detect a modest disease-independent decline in apparent fibrosis with increasing age (P<0.05), a finding we attribute to growth-associated muscle hypertrophy. Sirius red staining of endomysial connective tissue reflects fibre perimeters, while picric acid staining reflects myofibre cross-sectional area: the former thus increases linearly with growth, while the latter increases with the square of the diameter. Unlike mice (where substantial muscle growth has occurred by 6 weeks of age [42]), dogs are still growing between the 3-9-month timepoints used here [43] (the animals sampled here more than doubled in bodyweight over this time period), producing a downward shift in derived fibrosis fraction that our approach was able to measure.

**Figure 6:**
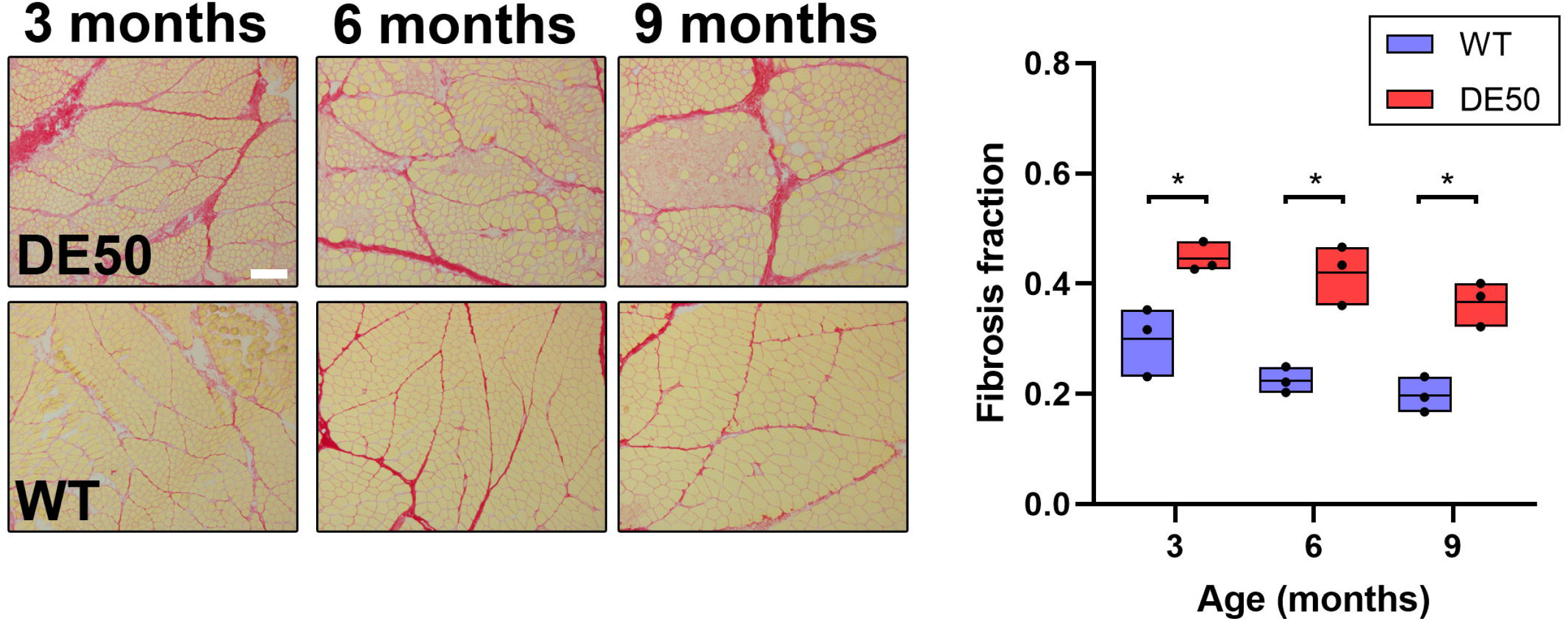
Fibrosis fraction quantification in healthy and dystrophic canine muscle Quantitative image analysis of vastus lateralis muscle biopsy samples collected from healthy (WT) and dystrophic (DE50) dogs at 3, 6, and 9 months of age reveals marked disease-associated differences (P<0.01, WT vs DE50-MD) and modest decreases in measured values with age (P<0.05, age). 2-way ANOVA with post-hoc multiple comparisons (Holm-Sidak) by age group as shown (* = P<0.05). Scalebar: 200µm.

### Rapid semi-automated measurement of muscle fibrosis

To streamline this analysis method, we developed a comparatively simple ImageJ macro (PSR_quantify.ijm [32]), capable of high-throughput analysis of large image collections (a step-by-step protocol is available at protocols.io [30]). As shown in figure 7, the user provides a folder of brightfield 10x images (in TIFF format) and a folder to place data (input/output directories, figure 7A), and supplies a unique analysis ‘sampleID’. The macro then extracts hue and saturation data from the entire image set and generates mean (+/-standard deviation) histograms for each (figure 7B, C), with thresholds for background subtraction (saturation) and yellow/red regions (hue) overlaid appropriately. Individual thresholds values can be adjusted and the overlay updated if necessary (figure 7D). Once ‘accept’ is selected, the macro then iterates through the image collection again (figure 7E). Hue and saturation channels are extracted from each image: saturation is used to generate a background mask, which is then applied to the hue channel. The final background-corrected hue image is used to extract yellow and red pixel counts and hence fibrosis fraction values. Any remaining pixels are counted as ‘other’. All values are then added to an excel-compatible comma-separated values (.csv) results table (figure 7F) for downstream analysis. For full transparency, fibrosis fraction values are exported both as “red vs yellow” (ignoring any pixels that fall under ‘other’) and as “total” (including those pixels), though these two derived values are essentially identical in most cases.

All image processing occurs in ‘batch mode’: no images are displayed. This makes analysis rapid (hundreds of images per minute) and presents the investigator with mean histograms rather than images, allowing thresholds to be selected without risk of bias. No source images are edited: all background subtraction and processing use temporary duplicates, allowing image collections to be archived, and analysis to be repeated with new parameters (if desired). The results table (figure 7F) stores all threshold values chosen within the column headers, and lists per-image fibrosis fractions, alongside counts of red, yellow, background and “other” pixels (any rare pixels not falling into the former three categories). All data (including raw histograms and derived mean and standard deviations) are automatically saved (as .csv) in the nominated folder, using a consistent nomenclature appended to the user-defined sampleID.

### HSB-based image segmentation is versatile

Having established the utility of HSB-based analysis for PSR staining, we investigated whether a similar approach could be extended to other colorimetric stains employed to assess muscle pathology. We selected acid phosphatase (AP), a brick-red stain for lysosomal enzyme activity, contrasted with a blue/purple haemalum counterstain. In dystrophic skeletal muscle this stain reveals degeneration/regeneration and macrophage infiltration, which is prominent in DE50-MD skeletal muscle [44]. As shown in figure 8, the same principal HSB-based methodology can be applied to this stain. Here segmentation via brightness eliminated slide background (figure 8A), while the red regions could be distinguished from counterstain using a combination of hue and saturation: hue defining the precise colour range, and saturation establishing a minimum threshold for stain intensity (figure 8B). This stain can thus also be quantified using a similar high-throughput methodology to that developed above, and accordingly we compiled a macro for this analysis (AP_quantify.ijm [33]). As shown in figure 8C, applying this approach to AP-stained canine muscle sections (as used for PSR analysis, above) revealed stark differences in staining between healthy and dystrophic muscles, with the focal nature of the pathology labelled by this stain (and concomitant image-to-image variation) suggesting a log distribution for quantitation. Analysis of our 3-9-month canine sample cohort (N=3, WT and DE50-MD, figure 8D) shows that, as with PSR, image-based quantitation is statistically tractable: HSB-based image segmentation thus represents a versatile approach to quantitative histology.

**Figure 7:**
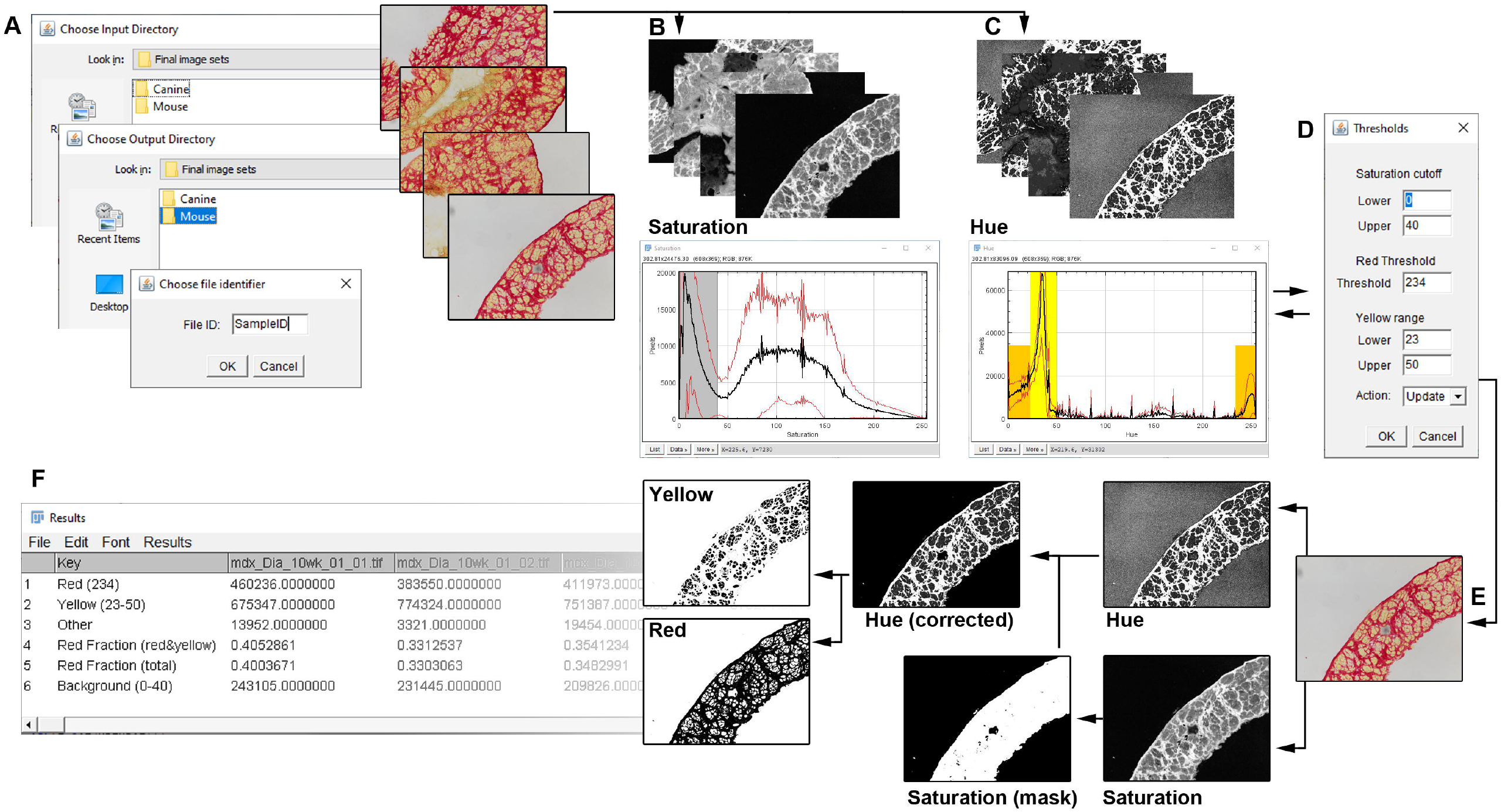
High-throughput semi-automated quantification of PSR stained muscle image collections using the HSB colour space Workflow for the PSR_quantify.ijm macro: running the macro (A) prompts the user to provide an input directory (where the images are located), an output directory (where data will be automatically saved as .csv files -this directory can be the same as the input directory), and a sampleID -the unique identifier that will be prefixed to all automatically saved data files. The macro then iterates through the entire collection, extracting saturation (B) and hue (C) histogram data for all images, which is then presented to the user as mean +/- standard deviation. Default thresholds for background subtraction (saturation data) and red/yellow brackets (hue data) are overlaid, and a menu provided to allow the investigator to adjust individual thresholds as necessary (D). Changing specific values and selecting ‘update’, then ‘ok’ changes the overlaid thresholds but does not alter the underlying data, and this can be repeated until the user is satisfied. The user then selects ‘accept’ and ‘ok’, and the macro iterates through the collection once more (E): for each image, the hue and saturation channels are extracted, with the saturation channel used to generate a background subtraction mask (based on selection background threshold) which is applied to the hue channel, from which red and yellow pixel counts can be extracted. Analysis is tabulated on a per-image basis (F), recording total counts of red and yellow pixels (and thresholds used to determine this), total counts of background (with threshold value used) and ‘other’ pixels (any image area that does not fall into any of the other categories), along with fibrosis fraction values, as “Red fraction (red vs yellow)” (ignoring ‘other’ pixels) and as “Red fraction (total)” (including all non-background pixels). Data is saved automatically in the output directory folder.

**Figure 8:**
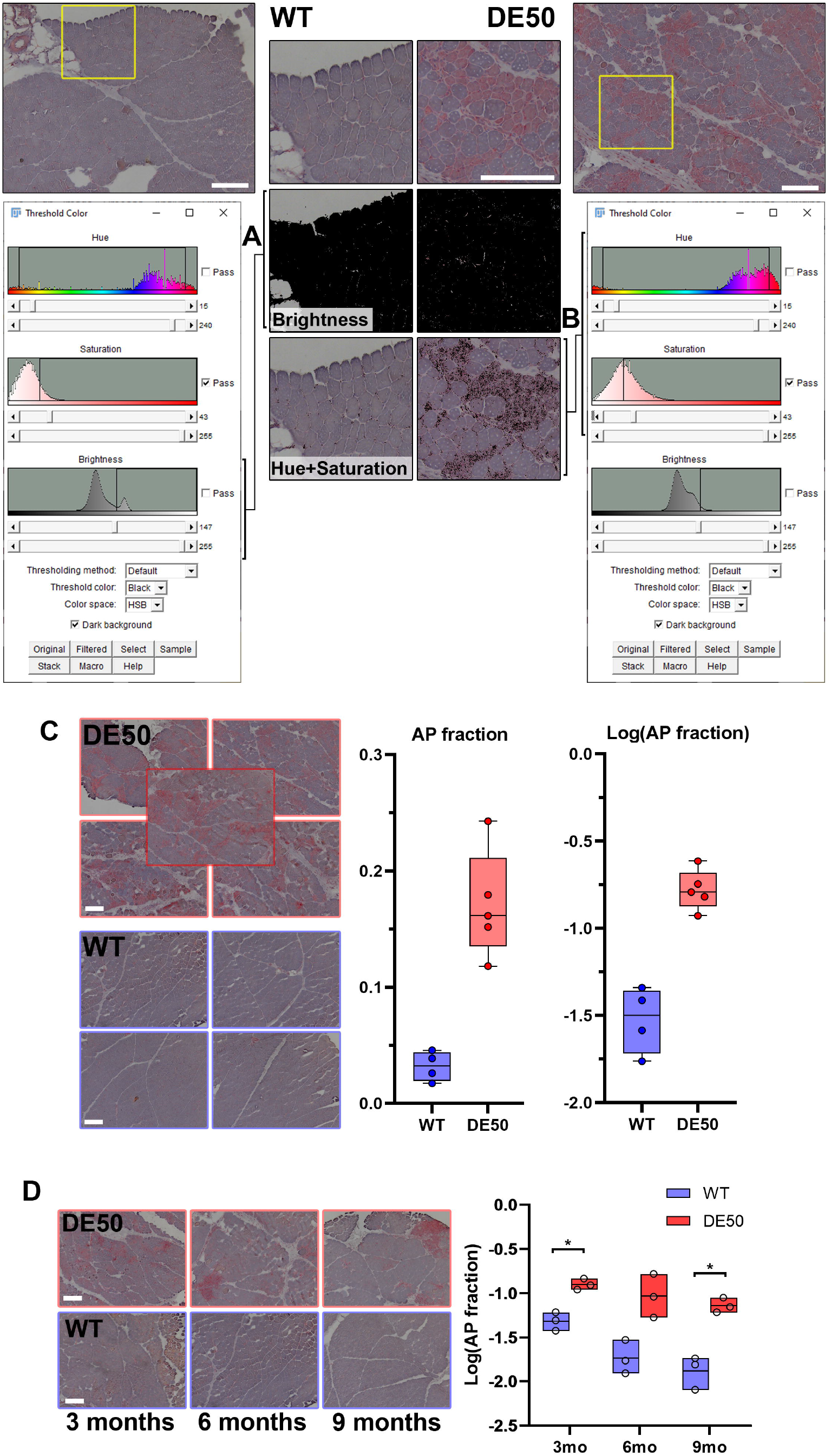
The HSB colour-space permits background elimination and stain segregation in acid phosphatase-stained muscle Canine dystrophic (DE50) *vastus lateralis* muscle shows widespread deposits of acid phosphatase activity, while healthy muscle (WT) does not. The HSB-based colour threshold tool in ImageJ permits background pixels to be eliminated via brightness gating (A), while hue can be used to designate red-stained pixels which are further restricted to a threshold stain intensity via saturation gating (B), and then expressed as a fraction of total (non-background) pixels: the ‘AP fraction’. As with PSR, the same parameters can be applied to both healthy and dystrophic muscle. This approach can be applied to multiple representative images DE50 and WT samples (C), allowing disease-specific increases in acid phosphatase activity to be quantified. Box and whisker plots show image-to-image variation is considerable, supporting a log distribution for this data: log(AP fraction). The AP_quantify.ijm macro allows mean log(AP fraction) values to be generated from *vastus* muscle biopsy samples collected from DE50 and WT dogs at 3, 6, and 9 months of age, revealing strong disease and age associated changes (P<0.01, DE50 vs WT, P<0.01, Age). 2-way ANOVA with post-hoc multiple comparisons (Holm-Sidak) by age group as shown (* = P<0.05). Boxes indicate mean and min/max values. Scalebars: 200µm.

## DISCUSSION

Quantifying fibrosis in muscle sections stained with picrosirius red is highly advantageous for monitoring disease progression and severity, and is essential for assessing therapeutic efficacy of interventions aimed at reducing or reversing this pathological feature. We present a fast and straightforward quantitation approach using the HSB colour space. Use of this assignment system separates image brightness and saturation from stain colour: this permits unbiased background subtraction on basis of saturation and renders analysis insensitive to any differences in illumination across the imaging field. Reducing colour to a single-dimension, intensity-independent, hue value moreover avoids the multi-dimensional complications inherent to other colour-spaces such as RGB or LAB. As shown here, hue maxima for red and yellow fractions are highly consistent between images, samples and staining batches, implying that these values are constants that reflect the innate chemistry of the dyes: barring minor differences in camera colour processing (for which we provide the option of adjusting thresholds accordingly), these values should be applicable to any muscle PSR staining, facilitating comparisons between studies, and between research groups and institutes. Segmentation by red/yellow fractions is simplistic to some extent: all intermediate shades are necessarily assigned to one or other fraction (there is no ‘orange’), but our approach is consequently intuitive, and allows a single quantitative metric to be derived (the ‘fibrosis fraction’), facilitating downstream analysis. Notably, we do not attempt to demark a threshold above which connective tissue represents ‘fibrosis’: the fibrosis fraction metric instead represents all connective tissue, whether heathy or pathological (circumventing the difficulties in delineating an arbitrary pathological boundary in the spectrum of fibrotic accumulation).

As demonstrated, this metric is also robust: the precise value of the red/yellow boundary is not critical, provided the same threshold is applied consistently. Per-image fibrosis fractions were remarkably consistent within samples, allowing both profound fibrotic changes and more subtle trends in fibrotic remodelling to be discerned and quantified.

The semi-automated open-source macro further developed here offers investigators a high-throughput means of deriving quantitative data from even substantial image collections. The onus remains with the investigator to capture representative images, but our macro allows subsequent analysis to be conducted swiftly, and with minimal risk of investigator bias. Selection of threshold values occurs before analysis, using mean histograms derived from the entire image set (blinding the investigator to individual sample identity), and analysis is rapid. Minimal image processing is required, and several hundred images can be measured in a matter of minutes, potentially affording investigations significant discriminatory power.

We further show that this HSB-based approach can be readily adapted to other colorimetric stains, using acid phosphatase as an example: as with PSR, this method allows quantitative data to be extracted from a representative sampling of images. AP staining is of limited use in the *mdx* mouse (where inflammatory muscle pathology is comparatively mild after the acute phase [34-36]), but this stain is valuable for more severely affected DMD models (such as the dog) and likely too for human pathology: as we demonstrate, robust quantification adds a valuable dimension, and renders this stain statistically tractable. It seems likely, given the data here, that other stains would be similarly amenable to quantitation. Oil red O (which labels lipid droplets vibrant red, alongside a blue nuclear counterstain) would be an obvious candidate, but this methodology could potentially be extended to more complex stains, such as trichrome.

## CONCLUSION

Quantitative analysis of colorimetric histological stains can reveal valuable biological information, but such analysis can also be challenging. Using the connective tissue stain picrosirus red, we show that use of the HSB colour space allows key quantitative parameters to be extracted from image data comparatively easily, and our data further suggests that this approach might well be adapted to additional histological stains.

## Supporting information

PSR_quantify macro

AP_quantify macro

