## Supplementary material for "Rapid histological quantification of muscle fibrosis and lysosomal activity using the HSB colour space": PSR_quantify macro: PSR_quantify.ijm

```
var lowerSat = 0;
var upperSat = 35;
var upperHue = 50;
var lowerHue = 21;
var LowerRed =234;


function processSaturation(input, output, filename)
{	
	open(input+filename);
	run("HSB Stack");
	run("Convert Stack to Images");
	nBins = 256;
	row = 0;
	selectWindow("Saturation");
	getHistogram(values, counts, nBins);
	
	//messy: don't want to store endless arrays in here, so write to the results then extract later
	for(i=0;i<nBins;i++)
	{
		setResult(filename, row, counts[i]);
		row++;
	}
	updateResults();
	close("*");
}

function processHue(input, output, filename)
{	
	open(input+filename);
	run("HSB Stack");
	run("Convert Stack to Images");
	nBins = 256;
	row = 0;
	selectWindow("Hue");
	getHistogram(values, counts, nBins);
	
	//messy: don't want to store endless arrays in here, so write to the results then extract later
	for(i=0;i<nBins;i++)
	{
		setResult(filename, row, counts[i]);
		row++;
	}
	updateResults();
	close("*");
}

function processSubtractedHue(input, output, filename) //for testing purposes
{	
	open(input+filename);
	run("HSB Stack");
	run("Convert Stack to Images");
	selectWindow("Saturation");
	setThreshold(lowerSat, upperSat); //gate off the low saturation
	run("Create Mask");
	imageCalculator("Add create", "Hue","mask");
	selectWindow("Result of Hue");

	nBins = 256;
	row = 0;
	getHistogram(values, counts, nBins);
	for(i=0;i<nBins;i++)
	{
		setResult(filename, row, counts[i]);
		row++;
	}
	updateResults();
	close("*");
}


function deriveStatistics(list)
{
	tempArray = newArray(list.length);
	meanArraySat = newArray(256);
	stDevArraySat = newArray(256);
	minArraySat = newArray(256);
	maxArraySat = newArray(256);
	meanArrayHue = newArray(256);
	stDevArrayHue = newArray(256);
	minArrayHue = newArray(256);
	maxArrayHue = newArray(256);
	j = 0;
	k = 0;
	mean = 0;
	selectWindow("SaturationData");
	IJ.renameResults("Results");
	for(j = 0; j< meanArraySat.length; j++)
	{
		for(i=0;i<list.length;i++)
		{
			if(endsWith(list[i],".tif"))
	    	{
				tempArray[i] = getResult(list[i],j); 
			}
		}
		Array.getStatistics(tempArray, min, max, mean, stdDev);
		meanArraySat[j] = mean;
		stDevArraySat[j] = stdDev; 
	}
	IJ.renameResults("SaturationData");
	run("Clear Results");
	for(i=0;i<meanArraySat.length;i++)
	{
			setResult(fileID+"Average",i,meanArraySat[i]);
			setResult(fileID+"StDev",i,stDevArraySat[i]);
			minArraySat[i] = meanArraySat[i] - stDevArraySat[i];
			maxArraySat[i] = meanArraySat[i] + stDevArraySat[i];
	}
	IJ.renameResults("SaturationMeanStDev");
	saveAs("Results",output+fileID+"SaturationMeanAndStDev.csv"); //export the analysis
	PlotSaturationGraph(meanArraySat,minArraySat, maxArraySat);
	run("Clear Results");

	for (i = 0; i<list.length;i++)
	{
		tempArray[i] = 0; //blank again
	}
	
	j = 0;
	k = 0;
	mean = 0;
	selectWindow("HueData");
	IJ.renameResults("Results");
	for(j = 0; j< meanArrayHue.length; j++)
	{
		for(i=0;i<list.length;i++)
		{
			if(endsWith(list[i],".tif"))
	    	{
				tempArray[i] = getResult(list[i],j);
			}
		}
		Array.getStatistics(tempArray, min, max, mean, stdDev);
		meanArrayHue[j] = mean;
		stDevArrayHue[j] = stdDev; 
	}
	IJ.renameResults("HueData");
	run("Clear Results");
	for(i=0;i<meanArrayHue.length;i++)
	{
			setResult(fileID+"Average",i,meanArrayHue[i]);
			setResult(fileID+"StDev",i,stDevArrayHue[i]);
			minArrayHue[i] = meanArrayHue[i] - stDevArrayHue[i];
			maxArrayHue[i] = meanArrayHue[i] + stDevArrayHue[i];
	}
	IJ.renameResults("HueMeanStDev");
	saveAs("Results",output+fileID+"HueMeanAndStDev.csv"); //export the analysis
	PlotHueGraph(meanArrayHue,minArrayHue, maxArrayHue);
	run("Clear Results");

	CheckParameters();
}

function CheckParameters() //here so we can loop back to this until happy
{
	Dialog.create("Thresholds"); //user specifies thresholds based on previous analysis
	Dialog.addMessage("Saturation cutoff");
	Dialog.addNumber("Lower",lowerSat);
	Dialog.addNumber("Upper",upperSat);
	Dialog.addMessage("Red Threshold");
	Dialog.addNumber("Threshold",LowerRed);
	Dialog.addMessage("Yellow range");
	Dialog.addNumber("Lower",lowerHue);
	Dialog.addNumber("Upper",upperHue);
	Dialog.addChoice("Action:", newArray("Update", "Accept", "Quit"));
	Dialog.show();
	
	str = Dialog.getChoice();
	if(str == "Accept")
	{
		lowerSat = Dialog.getNumber();
		upperSat = Dialog.getNumber();
		LowerRed = Dialog.getNumber();
		lowerHue = Dialog.getNumber();
		upperHue = Dialog.getNumber();
		close("*");
		AssessFibrosis();
	}
	if(str == "Update")
	{
		lowerSat = Dialog.getNumber();
		upperSat = Dialog.getNumber();
		LowerRed = Dialog.getNumber();
		lowerHue = Dialog.getNumber();
		upperHue = Dialog.getNumber();
		selectWindow("Saturation");
		close();
		selectWindow("Hue");
		close();
		PlotSaturationGraph(meanArraySat,minArraySat, maxArraySat);
		PlotHueGraph(meanArrayHue,minArrayHue, maxArrayHue);
		CheckParameters();
	}
	if(str == "Quit")
	{
		close("*");
		exit();
	}
}

function PlotSaturationGraph(mean, lower, upper)
{
	//plot the graph
	Plot.create("Saturation", "Saturation", "Pixels", mean);
	Plot.setLimitsToFit();
	Plot.getLimits(xMin, xMax, yMin, yMax);
	Plot.setColor("gray","lightGray");
	Plot.drawShapes("rectangles", lowerSat, yMax, upperSat, yMin); //this actually works! 
	Plot.setColor("red");
	Plot.add("line",lower);
	Plot.add("line",upper);	
	Plot.setColor("black");
	Plot.setLineWidth(2);
	Plot.setFrameSize(512, 304);
	Plot.show();
	selectWindow("Saturation");
	setBatchMode("show");
	setLocation(5, 5);
}

function PlotHueGraph(mean, lower, upper)
{
	//plot the graph
	Plot.create("Hue", "Hue", "Pixels", mean);
	Plot.setLimitsToFit();
	Plot.getLimits(xMin, xMax, yMin, yMax);
	Plot.setColor("yellow","yellow");
	Plot.drawShapes("rectangles", lowerHue, yMax, upperHue, yMin); 
	Plot.setColor("orange","orange");
	Plot.drawShapes("rectangles", LowerRed, yMax*0.5, xMax, yMin); 
	Plot.drawShapes("rectangles", xMin, yMax*0.5, lowerHue, yMin); 
	Plot.setColor("red");
	Plot.add("line",lower);
	Plot.add("line",upper);	
	Plot.setColor("black");
	Plot.setLineWidth(2);
	Plot.setFrameSize(512, 304);
	Plot.show();
	selectWindow("Hue");
	setBatchMode("show");
	setLocation(5, 464);
}

function AssessFibrosis()
{
	run("Clear Results"); //safety check
	setResult("Key", 0, "Red ("+LowerRed+")");
	setResult("Key", 1, "Yellow ("+lowerHue+"-"+upperHue+")");
	setResult("Key", 2, "Other");
	setResult("Key", 3, "Red Fraction (red&yellow)");
	setResult("Key", 4, "Red Fraction (total)");
	setResult("Key", 5, "Background ("+lowerSat+"-"+upperSat+")");
	updateResults();
	for (i = 0; i < list.length; i++)
	{	
		if(endsWith(list[i],".tif"))
	    {
	    	GetRedYellowBrackets(input, output, list[i]);
	    }
	}
	selectWindow("Results"); 
	saveAs("Results",output+fileID+"FibrosisFractions.csv"); //export the analysis
}

function GetRedYellowBrackets(input, output, filename)
{	
	open(input+filename);
	run("HSB Stack");
	run("Convert Stack to Images");
	selectWindow("Saturation");
	setThreshold(lowerSat, upperSat); //gate off the low saturation
	run("Create Mask");
	imageCalculator("Add create", "Hue","mask");
	selectWindow("Result of Hue");

	nBins = 256;
	row = 0;
	getHistogram(values, counts, nBins);
	
	//let's try and summarise the data
	totRed = 0;
	totYellow = 0;
	totOther = 0;
	totBackground = 0;
	RvsY = 0;
	RvsTotal = 0;
	
	for(i=0;i<nBins;i++)
	{
		if(i<(lowerHue -1)) //red
			totRed += counts[i];
		else if(i<upperHue) //yellow
			totYellow += counts[i];
		else if(i<(LowerRed+1)) //other 
			totOther += counts[i];
		else if(i>LowerRed && i<255) //more red: 255 is assigned the white masking, and is normally zero anyway
			totRed += counts[i];
		else if(i == 255)
			totBackground += counts[i];
	}
	
	RvsY = totRed/(totRed+totYellow);
	RvsTotal = totRed/(totRed+totYellow+totOther);
	
	setResult(filename, 0, totRed);
	setResult(filename, 1, totYellow);
	setResult(filename, 2, totOther);
	setResult(filename, 3, RvsY);
	setResult(filename, 4, RvsTotal);
	setResult(filename, 5, totBackground);
	updateResults();
	close("*");

}


input = getDirectory("Choose Input Directory");
output = getDirectory("Choose Output Directory");
Dialog.create("Choose file identifier");
Dialog.addString("File ID:", "")
Dialog.show();
fileID = Dialog.getString();
setBatchMode(true); 
list = getFileList(input);
run("Clear Results"); //safety check
for (i = 0; i < list.length; i++)
{	
	if(endsWith(list[i],".tif") || endsWith(list[i],".TIF"))
    {
    	processSaturation(input, output, list[i]);
    }
}
selectWindow("Results");
saveAs("Results",output+fileID+"SaturationHistograms.csv"); //export the analysis
IJ.renameResults("SaturationData");
run("Clear Results"); //safety check
for (i = 0; i < list.length; i++)
{	
	if(endsWith(list[i],".tif") || endsWith(list[i],".TIF"))
    {
    	processHue(input, output, list[i]);
    }
}
selectWindow("Results");
saveAs("Results",output+fileID+"HueHistograms.csv"); //export the analysis
IJ.renameResults("HueData");
run("Clear Results"); //safety check
for (i = 0; i < list.length; i++)
{	
	if(endsWith(list[i],".tif") || endsWith(list[i],".TIF"))
    {
		processSubtractedHue(input, output, list[i]);
    }
}
selectWindow("Results");
saveAs("Results",output+fileID+"SubtractedHueHistograms.csv"); //export the analysis
run("Clear Results"); //safety check
//then process the results
deriveStatistics(list);


setBatchMode(false);
```
