## Supplementary material for "Rapid histological quantification of muscle fibrosis and lysosomal activity using the HSB colour space": AP_quantify macro: AP_quantify.ijm

```
var lowerSat;
var upperSat;
var upperHue;
var lowerHue;
var saturation;

//gate off brightness to exclude background, then restrict acid phos only to red of a given saturation


function GetAcidPhos(input, output, filename)
{	
	open(input+filename);
	run("HSB Stack");
	run("Convert Stack to Images");
	selectWindow("Brightness");
	setThreshold(lowerSat, upperSat); //gate off the high brightness
	run("Create Mask");
	imageCalculator("Add create", "Hue","mask");
	selectWindow("mask");
	close();
/*	selectWindow("Saturation");
	nBins = 256;
	getHistogram(values, counts, nBins);
	//have a rough stab at the peak saturation
	maxArray = newArray(3);
	valArray = newArray(3);

	//be cheap, screw it
	for(j=0;j<3;j++)
	{
		for (i = 0; i < nBins; i++) 
		{
			if(counts[i] > maxArray[j])
			{
				maxArray[j] = counts[i];
				valArray[j] = i;
			}
		}
		int = valArray[j];
		counts[int] = 0; //blank the highest
	}
	//get average
	int = 0;
	for(j=0;j<3;j++)
	{
		int += valArray[j];
	}
	int = int/3;
	int+=10;*/
	selectWindow("Result of Hue");
	//get literally ALL the non-background pixels
	nBins = 256;
	totCount = 0;
	totBackground = 0;
	getHistogram(values, counts, nBins);
	for(i=0;i<nBins;i++)
	{
		if(i < 255)
			totCount += counts[i];
		else 
			totBackground += counts[i];
	}

	//now mask off all but high saturation red
	selectWindow("Saturation");
	setThreshold(saturation, 255); //gate off the low saturation
	run("Create Mask");
	run("Invert");
	imageCalculator("Add create", "Result of Hue","mask");
	selectWindow("Result of Result of Hue");
	nBins = 256;
	getHistogram(values, counts, nBins);

	totRed = 0;

	for(i=0;i<nBins;i++)
	{
		if(i<(lowerHue -1)) //red
			totRed += counts[i];
		else if(i>=upperHue && i<255) //red: 255 is assigned the white masking, and is normally zero anyway
			totRed += counts[i];
	}
	RedFraction = totRed/(totCount);
	
	setResult(filename, 0, totRed);
	setResult(filename, 1, totCount);
	setResult(filename, 2, RedFraction);
	setResult(filename, 3, totBackground);
	updateResults();
	close("*");

}


input = getDirectory("Choose Input Directory");
output = getDirectory("Choose Output Directory");
Dialog.create("Choose file identifier");
Dialog.addString("File ID:", "")
Dialog.show();
fileID = Dialog.getString();
setBatchMode(true); 
Dialog.create("Bright/Sat Thresholds"); //user specifies thresholds based on previous analysis
Dialog.addNumber("LowerB",200);
Dialog.addNumber("UpperB",255);
Dialog.addNumber("Saturation", 43);
Dialog.show();
lowerSat = Dialog.getNumber();
upperSat = Dialog.getNumber();
saturation = Dialog.getNumber(); 
Dialog.create("Red Boundaries"); //user specifies thresholds based on previous analysis
Dialog.addNumber("Lower",15);
Dialog.addNumber("Upper",240);
Dialog.show();
lowerHue = Dialog.getNumber();
upperHue = Dialog.getNumber();
//Dialog.create("Yellow Threshold"); //user specifies thresholds based on previous analysis
//Dialog.addNumber("Lower",21);
//Dialog.addNumber("Upper",41);
//Dialog.show();
//lowerHue = Dialog.getNumber();
//upperHue = Dialog.getNumber();
list = getFileList(input);
//label the results table
setResult("Key", 0, "Red ("+lowerHue+"-"+upperHue+")");
//setResult("Key", 1, "Yellow ("+lowerHue+"-"+upperHue+")");
setResult("Key", 1, "Total ("+saturation);
setResult("Key", 2, "Red Fraction");
//setResult("Key", 4, "Red Fraction (total)");
setResult("Key", 3, "Background ("+lowerSat+"-"+upperSat+")");
updateResults();
for (i = 0; i < list.length; i++)
{	
	if(endsWith(list[i],".tif"))
    {
    	GetAcidPhos(input, output, list[i]);
    }
}
selectWindow("Results"); 
saveAs("Results",output+fileID+"AcidPhosFractions.csv"); //export the analysis

setBatchMode(false);
```
